# Detecting rare copy number variants (CNVs) from Illumina genotyping arrays with the CamCNV pipeline: segmentation of z-scores improves detection and reliability

**DOI:** 10.1101/2020.04.23.057158

**Authors:** Joe Dennis, Logan Walker, Jonathan Tyrer, Kyriaki Michailidou, Douglas F. Easton

## Abstract

**Background:** The intensities from genotyping array data can be used to detect CNVs but a high level of noise in the data and overlap between different copy-number intensity distributions produces unreliable calls, particularly when only a few probes are covered by the CNV.

**Results:** We present a novel pipeline (CamCNV) with a series of steps to reduce noise and detect more reliably rare CNVs covering as few as three probes. The method uses the information from all samples to convert intensities to z-scores, thus adjusting for variance between probes. We tested the sensitivity of our pipeline by looking for known CNVs from the 1000 Genomes project in our genotyping of 1000 Genomes samples. We also compared the CNV calls for 1,661 pairs of genotyped replicate samples. At the chosen mean z-score cut-off, sensitivity to detect the 1000 Genomes CNVs was approximately 85% for deletions and 65% for duplications. From the replicates we estimate the false discovery rate is controlled at ∼10% for deletions (falling to below 3% with more than five probes) and ∼28% for duplications. The pipeline demonstrates improved sensitivity when compared to calling with PennCNV, particularly for short deletions covering only a few probes

**Conclusion:** The CamCNV pipeline provides a reliable method of detecting rare CNVs from Illumina array data and can be used for CNVs that only cover a few probes. For each called CNV the mean z-score is a useful metric for controlling the false discovery rate.

## 1 Introduction

Genotyping experiments with large case control studies have been effective at identifying SNPs associated with disease. Associations with common CNVs can also be detected by imputation of their genotypes using reference datasets with sequence data. The 1000 Genomes Project reported that 73% of structural variants with >1% frequency are in strong linkage disequilibrium (*r*^2^ > 0.6) with nearby SNPs, although the proportion falls to 44% for common duplications (1). For example, at two GWAS loci associated with breast cancer, the APOBEC locus (2, 3) and 2q35 (4), deletions tagged by SNPs are associated with risk, and the CNV has been identified as the probable causal variant in each case. However, with current imputation reference panel sizes it is not possible to reliably impute rare CNVs. With increasing sample sizes genotyped on arrays there is an opportunity to detect novel associations with rare CNVs that potentially have large effects on disease risk.

Disease associations for rare CNVs detected using genotyping arrays have typically involved large CNVs, often covering multiple genes, for example those associated with developmental disorders (5) and schizophrenia (6). Evidence from clinical tests using gene panels suggests that intragenic CNVs account for a significant proportion of pathogenic variants, between 4.7% to 35% depending on the clinical area.(7) There is also evidence that intronic CNVs can affect the regulation of genes where the exons are highly conserved across evolution. (8) Such CNVs are likely to cover only a small number of probes on genotyping arrays.

Several methods have been developed for calling CNVs from the intensities of genotyping arrays. A commonly used method is PennCNV (9), which implements a Hidden Markov Model to find increases or decreases in the intensities of probes, ordered by genomic position, for a single sample. Other methods such as DNAcopy (10) use circular binary segmentation to detect these shifts in intensity by sample. Methods such as CNVTools (11) and cnvHap (12) model the intensities from all samples at CNV loci but are designed to analyse more common CNVs.

The pipeline (CamCNV) (Figure 1) we describe here uses DNAcopy but also aims to incorporate the information from all samples. It works on a similar principle to the MeZOD method (13, 14) to detect rare outliers in the distribution of intensities. The method excludes probes in common CNV regions and assumes that at each probe tested the vast majority of samples will have two-copies of DNA and their intensities will form a normal distribution. Single copy deletions or three-copy duplications will form part of separate distributions that overlap with the two-copy distribution. Converting the intensity measurements to z-scores and segmenting using DNAcopy detects runs of probes that are potential CNVs for each sample. Our method draws on previous work (15) to remove noise and batch effects from the intensities with a principal component adjustment. In addition it uses the information from across each sample to exclude potential CNVs where the shift in intensity does not correlate with the shift for known CNVs. We provide estimates for the sensitivity using known CNVs in 1000 Genomes samples and for the false discovery rate from a large number of replicates.

**Figure 1.**
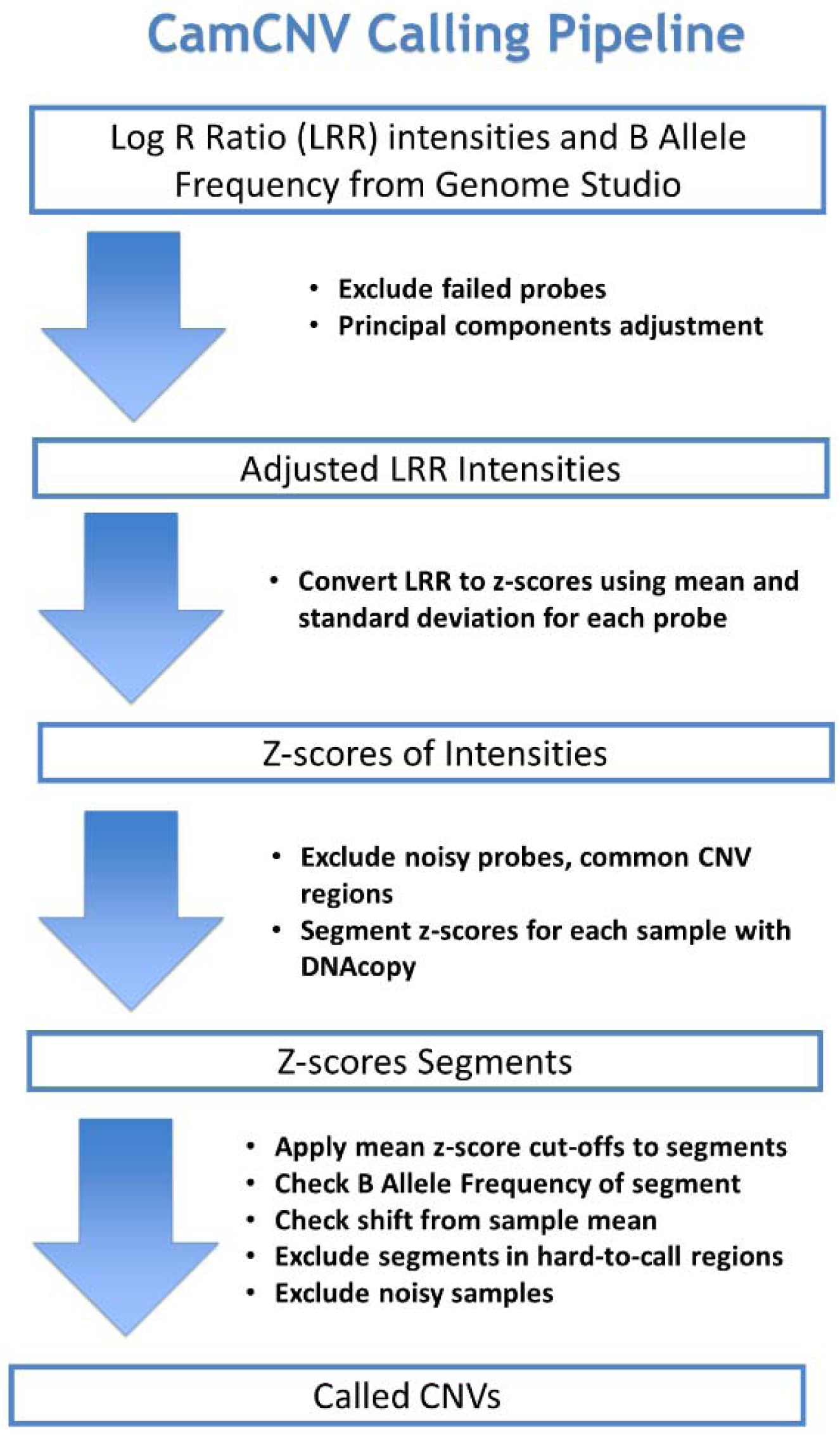
CNV calling pipeline (CamCNV)

## 2 Methods / Pipeline

### 2.1 Principal component adjustment

We first export the Log R Ratio (LRR) intensities for each sample at each probe from Illumina’s Genome Studio software into a matrix ordered by probe position. In order to reduce batch effects in the intensity measurements, we follow the advice of Cooper et al. (15) and perform a principal component adjustment (PCA) on the LRR. This should also adjust for genomic waves. (16) We exclude from the matrix SNPs that fail genotyping quality control and use the “thin” function from the R bigpca package (17) to select 15% of SNPs most representative of the principal components. We use the function “quick.elbow” to decide how many principal components lie above the “elbow” of the PCA plot and adjust the entire LRR matrix for all SNPs for those PCs using the bigpca PC.correct function.

### 2.2 Z-score calculation

Using all samples that pass QC, we calculate the means and standard deviations for the LRR at each probe and hence convert the LRRs to z-scores as:

(Sample LRR – Mean LRR for probe)/Standard Deviation of LRR for probe.

### 2.3 Probe and loci exclusions

Before calling CNVs we then exclude probes likely to produce unreliable calls. We exclude low intensity probes (Illumina metric AB R Mean <0.2) and those not clustered by the Illumina Gentrain algorithm, which are likely to represent technical failures at the chip genotyping level. As our pipeline is designed to detect rare CNVs, we exclude probes within CNVs reported by the 1000 Genomes project as having greater than 1% frequency. We observed some false positive deletion calls at probes within a few base pairs of another probe, presumably because competition for DNA fragments lowers the intensity. To guard against this, we exclude probes within three base pairs of another probe. Finally, we exclude noisier probes with high LRR variance (choosing a threshold of 0.12 - two standard deviations above the mean variance for all probes.) After CNV calling, we exclude CNVs falling within 1MB of the telomeres and centromeres as defined in the “gap” database table from the UCSC Genome Browser. We also exclude CNVs within immune-related loci (MHC, T-Cell receptor and immunoglobulin heavy chain), as CNVs at these loci may occur somatically in blood.

### 2.4 Circular binary segmentation (CBS) of z-scores

The z-scores are saved into an R data frame for each chromosome ordered by genomic position with the excluded probes removed. We use the R package DNACopy (10, 18) to segment the z-scores for each sample individually by chromosome using circular binary segmentation. After using the ‘smooth’ function to detect outliers and smooth the data, we call the segments using the segment function. This produces a list of segments for each sample for each chromosome with the start and end positions, number of probes and the segmentation mean for each segment. Before deciding the final cut-off values we label as potential CNVs segments with a mean z-score below −2 or above +1.5, covering between three and 200 probes.

### 2.5 LRR shift scores per sample

For each sample we calculate the mean LRR for that sample for that chromosome. To ensure that this measure is based on normal two-copy intensities, we only use common SNPs with a high call rate (>99%) where the sample appears heterozygous (BAF around 0.5). For each potential CNV we calculate the difference between the mean sample LRR for the chromosome and the mean LRR for the CNV. Based on the shifts for known CNVs, we exclude CNVs where this difference lies outside the range of 0.2 and 0.8 for deletions and 0.1 and 0.4 for duplications. For these calculations we use the LRR before the principal component adjustment.

### 2.6 B Allele Frequency (BAF) score

The B Allele Frequency can be used alongside the LRR to identify CNVs. For samples with normal two-copy DNA at a probe, this score should be near zero for homozygotes for the reference allele (AA), 0.5 for heterozygotes (AB) and one for homozygotes for the alternate allele (BB). For deletions, the BAF has some value as a negative predictor; any values not close to zero (A-) or one (B-) suggest that it is not a true deletion. Thus, for each potential deletion (as an exclusion score) we calculate the percentage of probes with BAF > 0.3 and <0.7. We exclude deletions in which ≥25% of probes fail the BAF check.

For three copy duplications, scores should be around zero (AAA), one (BBB), 0.33 (AAB) or 0.66 (ABB), so we can record a positive predictor score as the percentage of probes with BAF >0.1 and <0.4 or >0.6 and <0.9. However for short CNVs the value of the BAF is limited as we do not necessarily expect the CNV to cover probes that might be polymorphic, particularly on arrays targeting rare variants.

## 3 Test data

### 3.1 BCAC samples

To develop the pipeline we used data from 49,923 breast cancer cases and 42,148 controls of European ancestry from 54 studies in the Breast Cancer Association Consortium (BCAC) dataset. These were genotyped on the Illumina OncoArray chip (533,631 probes) at five genotyping centres. (19). We included 1,661 pairs of duplicate samples. While the majority of duplicates were genotyped intentionally by studies for quality control, 160 of the pairs were submitted by different studies and later identified as the same person from their genotypes and date of birth. The principal component adjustment described above was performed separately on each study. For eight studies the number of samples was too large to adjust as a single batch and these studies were divided into two or three batches.

### 3.2 1000 Genomes Project controls

As part of the OncoArray genotyping quality control, samples from the 1000 Genomes project were added to each plate at the genotyping centres. In addition, 240 samples were genotyped by Illumina. After sample quality control we used OncoArray data for 233 individuals comprising 2,226 replicates. 169 individuals were genotyped once and 64 were genotyped more than once (between two and 2 and 349 times). We downloaded the structural variant calls from Phase3 of the 1000 Genomes project (1), extracted bi-allelic deletions and duplications for the 233 comparison individuals and calculated the start and end probes for any variant covered by OncoArray probes. We excluded large variants covering more than 50 probes. There are some complex variants (for example one individual has a series of 60 deletions at the end of chromosome 18). To simplify the test we excluded sets of CNVs where a sample had five or more CNVs starting in the same megabase.

### 3.3 Sample quality control

For the BCAC dataset we included only DNA samples extracted from whole blood to minimise variation in sample quality. 1000 Genomes DNA samples were from lymphoblastic cell lines (LCLs). Samples with genotype call rate below 99% or excess heterozygosity or sex chromosome anomalies were excluded. (2) A derivative log ratio spread (DLRS) figure was calculated for each sample as the standard deviation of the differences between LRR for probes ordered by genomic position, divided by the square root of two. This measures the variance from each probe to the next averaged over the whole genome and so it is insensitive to fluctuations in intensities between different chromosomes for individual samples. (15) We excluded samples with DLRS 3.5 standard deviations above the DLRS mean for the study. This exclusion was applied both before and after the principal component adjustment.

After the segmentation we counted the number of short segments (between three and 200 probes) for each sample. We observed that the distribution of segment counts is skewed with many samples with an excessive of number of segments. As an exclusion applied to all samples, we calculated a cut-off for the maximum number of segments using the following formula where x is the segment count for each sample (based on the rationale that the true number of segments should be approximately Poisson):

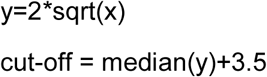

### 3.4 Imputed CNVs

BCAC genotypes were phased using ShapeIT and imputed to the 1000 Genomes Phase 3 reference panel using Impute2 as previously described (2). We compared called CNVs for samples with imputed dosages of one i.e. those imputed to have one or three copies at the CNV locus.

### 3.5 PennCNV calling

As a comparison we also called all samples using PennCNV with the LRR adjusted for principal components as the input. GC model files were calculated for the OncoArray giving the percentage of G or C bases for 500 Kb either side of each probe. Population-specific files were calculated to give the frequency of the B allele for each SNP. After calling, the PennCNV script clean_cnv.pl was used to merge neighbouring CNVs where the gap between them was less than 20% of the total length of the combined CNVs. The PennCNV script filter_cnv.pl was used to generate quality control statistics for each sample. Samples with LRR Standard Deviation > 0.2 or an excessive number of CNVs were excluded.

### 3.6 Comparison of CNV calls

In the comparisons below, CNV calls were counted as an exact match if they covered the maximum number of probes on OncoArray for the reference CNV and were no longer. We categorised calls as partial matches if at least 50% of the expected probes were covered and/or where the called CNV extended further, excluding those covering greater than five times the number of probes in the reference CNV.

## 4 Results

### 4.1 Distribution of mean z-scores for known CNVs

To evaluate the distribution of mean z-scores for segments matching known CNVs, we compared the segments called by the pipeline that matched a set of well-imputed rare CNVs in the BCAC data. We identified 26 distinct biallelic deletions and 11 biallelic duplications, carried by 7,189 and 4,874 samples, respectively. We plotted histograms of the mean z-scores of the segments that perfectly matched the imputed CNVs (n=4,881 deletions and n=1,051 duplications) as shown in Figure 2.

**Figure 2:**
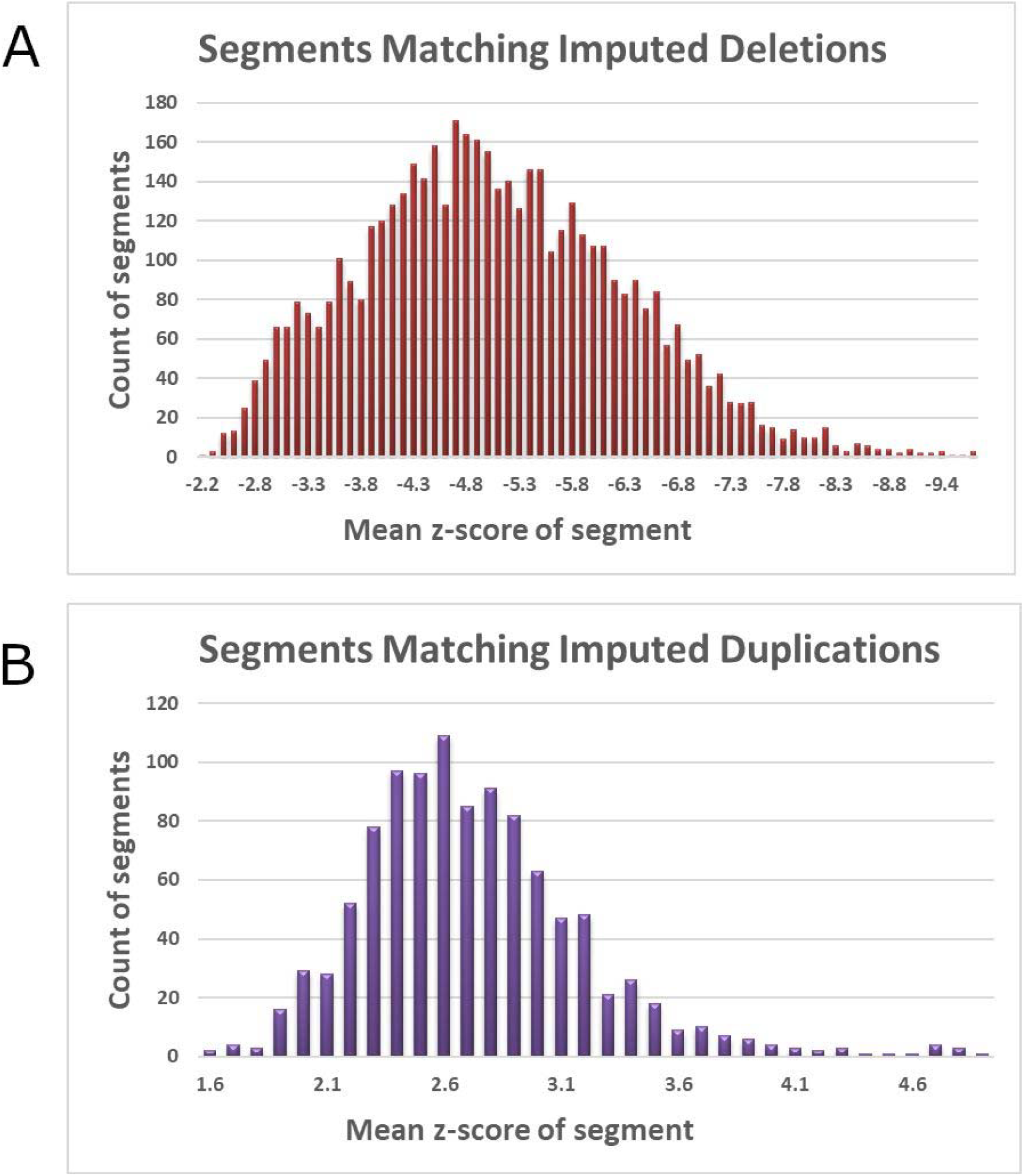
Mean z-score of intensity of segments matching well-imputed CNVs in the BCAC data for A) 26 deletions, B) 11 duplications.

The overall match rate for imputed data using our chosen cut-offs (see below) is summarised in Table 1. The match rates for these CNVs will be affected by the uneven distribution of probe coverage (Table 2). The majority of these deletions only have three probes while most of the duplications have more than six.

**Table 1.** Match rate for called CNVs against BCAC imputed CNVs

|  | Separate CNVs | Samples | Matched Rate | Perfect Matches | Matches excluded by QC |
| --- | --- | --- | --- | --- | --- |
| Deletions | 26 | 7189 | 77.9% | 65.2% | 4.8% |
| Duplications | 11 | 4874 | 61.1% | 20.7% | 5.9% |

**Table 2.** Probe coverage for CNVs in comparison with imputed data

| Probes Covered | Deletions | Duplications |
| --- | --- | --- |
| 3 | 14 | 1 |
| 4 | 3 | 2 |
| 5 | 3 | 1 |
| 6 to 10 | 3 | 5 |
| >10 | 3 | 3 |

### 4.2 Z-score cut-offs and estimated false discovery rate (FDR)

To select mean z-score cut-offs to control the false discovery rate, we compared called CNVs in 1,661 pairs of BCAC replicates. In doing this, we make the assumption that a CNV seen in both replicates is likely to be genuine, whereas a CNV seen in only one sample is likely to be an artefact. This is reasonable if the calling errors in the replicates are truly independent, but it is possible that the errors are correlated due to the local sequence or other issues related to the individual samples; hence this approach gives a lower bound on the FDR.

We observed that the mean z-scores of the unmatched calls from the first of the replicate pairs (Figure 3) are concentrated towards the normal end of the distribution. This suggests that they are mostly false positive calls at loci with normal two-copy DNA, but it is also possible that some are genuine CNVs at probes with a large degree of overlap with the normal distribution (hence harder to call). The relationship between the mean z-score of the segment and the proportion that are unmatched is shown in Figure 3. To provide a reasonable control of the FDR, while not impacting the sensitivity too severely, we chose cut-offs at −3.7 for deletions and +2 for duplications; this results in an overall FDR of ∼10% for the remaining deletions and ∼30% for duplications (Figure 4). The FDR does not reduce materially for duplications with more stringent cut-offs. At the other end of the distributions, we observed a small number of extreme values for the mean z-score. These may reflect homozygous deletions or duplications with more than three copies but could also be due to probes failing to bind at all. We therefore excluded calls with z< −14 for deletions and z> +10 for duplications. While this could exclude some true homozygotes, the main focus is on calling call rare CNVs and hence this possibility can be ignored here.

**Figure 3.**
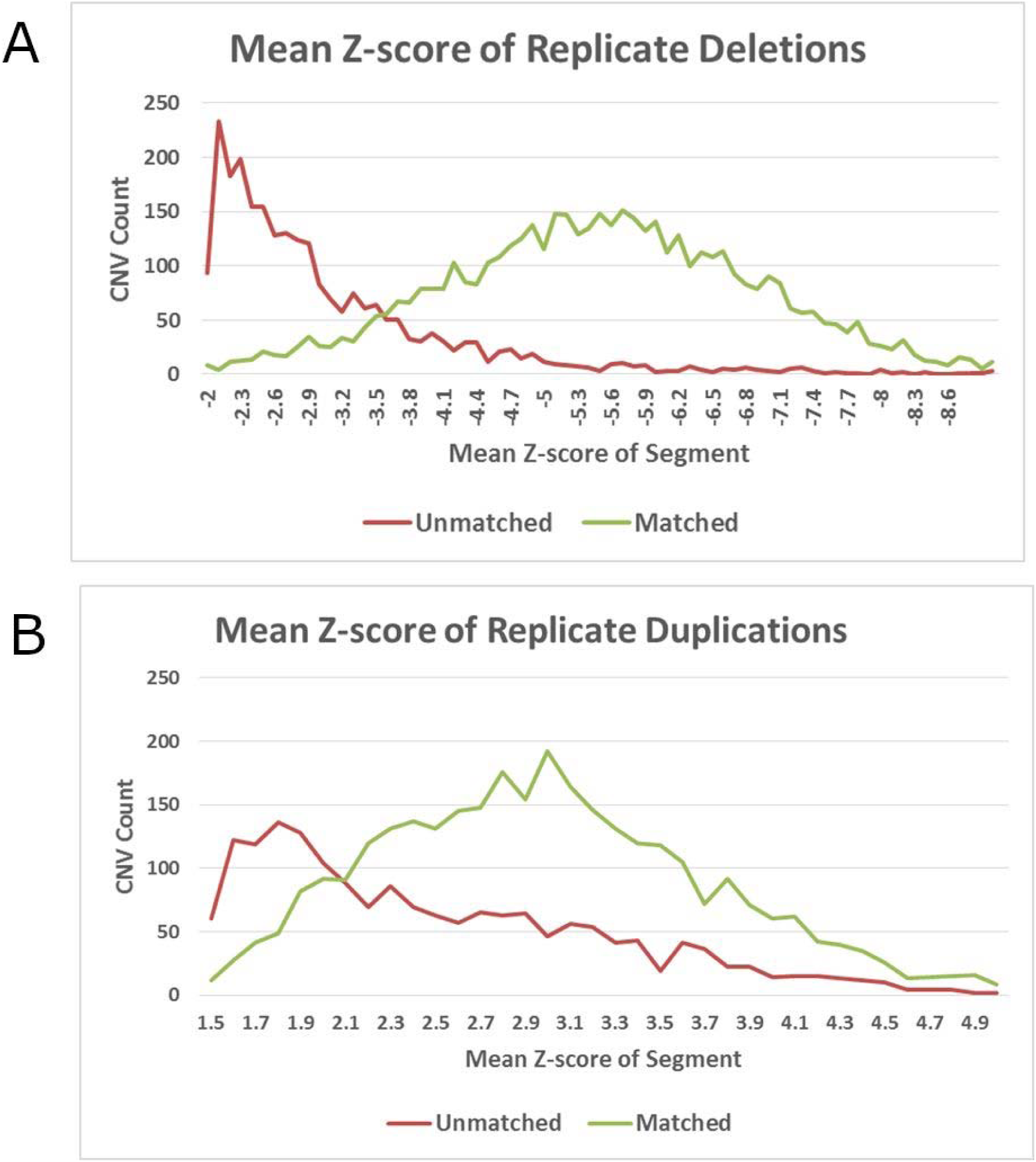
Mean z-scores of matching and umatching deletions (A) and duplications (B) in replicate samples

**Figure 4.**
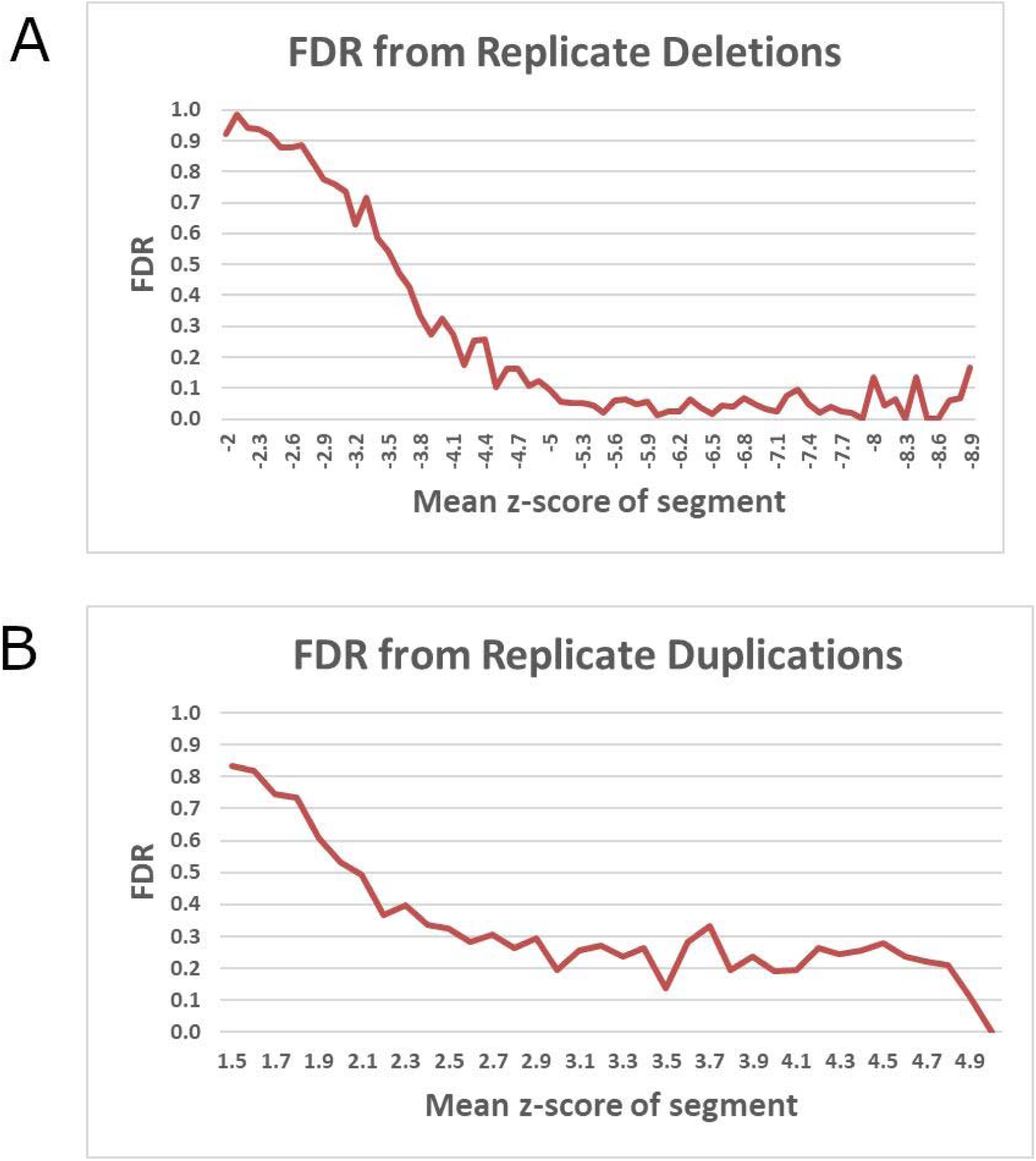
Estimated false discovery rate by mean z-score of deletions (A) and duplications (B) in replicate samples

Using these cut-offs, the results from the matching exercise are shown in Table 3. Based on exact or partial matches, the overall FDR was 10% for deletions and 28% for duplications. The FDR decreases with increasing numbers of probes as shown in Table 4, particularly for deletions. We observed no significant difference between the unmatched rate for samples from the same study and those collected by different studies (n=160 pairs) where the DNA has been extracted by different laboratories at different ages and one might expect more variability from batch effects.

**Table 3.**
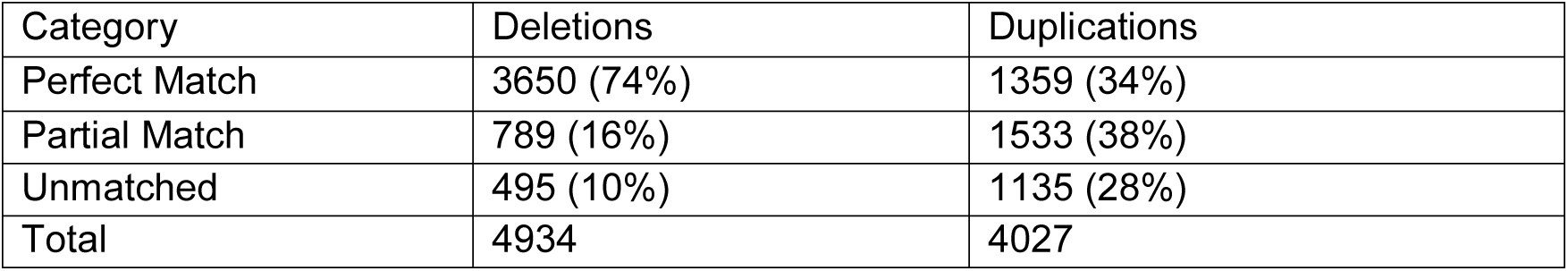
Match rate for CNV calls from 1,661 pairs of replicate samples in BCAC data

**Table 4.** False discovery rate by probe length from comparison of replicate sample calls

| Probes | Deletions |  | Duplications |  |
| --- | --- | --- | --- | --- |
|  | Count | FDR | Count | FDR |
| 3 | 1541 | 23.4% | 489 | 62.4% |
| 4 | 852 | 8.9% | 520 | 43.1% |
| 5 | 479 | 5.8% | 382 | 43.7% |
| 6 | 288 | 2.4% | 282 | 36.5% |
| 7 | 231 | 2.2% | 259 | 40.5% |
| 8 | 172 | 1.7% | 223 | 19.3% |
| 9 | 170 | 0.6% | 193 | 23.3% |
| 10+ | 1201 | 1.2% | 1679 | 8.5% |

### 4.3 Summary of called CNVs

Using the above cut-offs we called a mean of 2.9 deletions (SD 1.6) and 2.5 duplications (SD 2.0) per sample among 92,071 samples in the BCAC dataset. Relaxing the cut-offs to −2 for deletions and 1.5 for duplications would give 5.4 deletions (SD 5.2) and 3.4 duplications (SD 2.7) per sample. After excluding long CNVs covering more than 50 probes, the mean length was 7.3 probes (SD 7.1) / 34 kilobases (kb) (SD 71 kb) for deletions, and 10.0 probes (SD 9.6) / 70 kb (SD 109 kb) for duplications. 14% of the probes on the chip were covered by five or more deletions and 22% of probes by more than five duplications. As designed, our pipeline did not detect the more common CNVs. The probes towards the upper end of our frequency range showed a large degree of overlap with previously published CNV regions. For example 58% of probes in deletions and 39% of probes in duplications with frequency >0.5%, fall within a map of CNV regions compiled from high-confidence entries in the Database of Genomic Variants (DGV) (20). (Supplementary Table 1)

### 4.4 Detection of known CNVs for 1000 Genomes samples

We compared our pipeline calls for 1000 Genomes samples genotyped on the OncoArray with the CNVs published by the 1000 Genome project. We identified 523 rare (<1% frequency) bi-allelic deletions and 308 rare duplications that cover at least three probes on OncoArray. Since some CNVs were carried by more than one of our comparison samples and some samples were replicated multiple times in the OncoArray dataset (n=3,532 comparison CNV calls in total), we chose a random replicate to perform the comparisons. The results are shown in Table 6. 80.2% of deletions and 59.5% of duplications were detected with either a perfect or partial match. CNVs that were not detected at all tended to be short (60% cover only three or four probes) and tended to have a smaller proportion covered by the chip probes. Those detected with 100% precision have a mean 65% (SD 22%) of base pairs covered compared to a mean of 53% (SD 25%) for CNVs with no matches. Around 6% of CNVs were detected but were excluded as they were outside the mean z-score cut-offs or failed other QC metrics. Where CNVs were carried by more than three replicates, deletions called in one sample tended to be called in the majority of samples, with a poorer replication rate for duplications. 70% of these detected deletions (n=63) and 40% of duplications (n=35) had a greater than 90% replication rate. (Supplementary Figure 1)

**Table 6.** Comparison of called CNVs for 1000 Genomes samples with CNVs from sequencing

| Category | Deletions | Duplications |
| --- | --- | --- |
| Perfect match | 253 (48.2%) | 43 (14%) |
| Partial match | 168 (32%) | 140 (45.5%) |
| Matched but outside mean z-score cut-offs | 27 (5.1%) | 11 (3.6%) |
| Matched but excluded by QC metrics | 6 (1.1%) | 3 (1%) |
| Called CNV >5x probes of 1KG CNV | 6 (1.1%) | 7 (2.3%) |
| Uncalled | 65 (12.4%) | 104 (33.8%) |

### 4.5 Comparison with PennCNV calling

We also called the 1000 Genomes samples and BCAC replicates using PennCNV. For the 1000 Genomes comparison, we categorised the calling of the 525 deletions and 308 duplications as either “perfect”, if the CNV call matched exactly with the known genotype in more than 90% of replicates, and “imprecise” if at least some calls partially matched or the replicate was below 90%. Table 7 and Figure 5 show the comparison between PennCNV and CamCNV calling. Overall CamCNV called 20% more deletions and 10% more duplications than PennCNV. The set of CNVs (Figure 6) not detected by PennCNV tended to cover fewer probes: the deletions covered a mean of 3.9 probes compared to 8.5 probes for those also matched by PennCNV, while CamCNV covered 9.6 probes for duplications compared to 13.8 probes for PennCNV.

**Table 7.** Comparison of sensitivity of CamCNV and PennCNV calling of 1000 Genomes CNVs

| Deletions |  | PennCNV calling |  |  |
| --- | --- | --- | --- | --- |
|  |  | Perfect Match | Imprecise Match | No matches |
| CamCNV Calling | Perfect Match | 24% | 7% | 11% |
|  | Imprecise Match | 3% | 29% | 11% |
|  | No matches | 1% | 1% | 12% |

| Duplications |  | PennCNV calling |  |  |
| --- | --- | --- | --- | --- |
|  |  | Perfect Match | Imprecise Match | No matches |
| CamCNV Calling | Perfect match | 6% | 5% | 3% |
|  | Imprecise Match | 2% | 42% | 15% |
|  | No matches | 1% | 7% | 21% |

**Figure 5.**
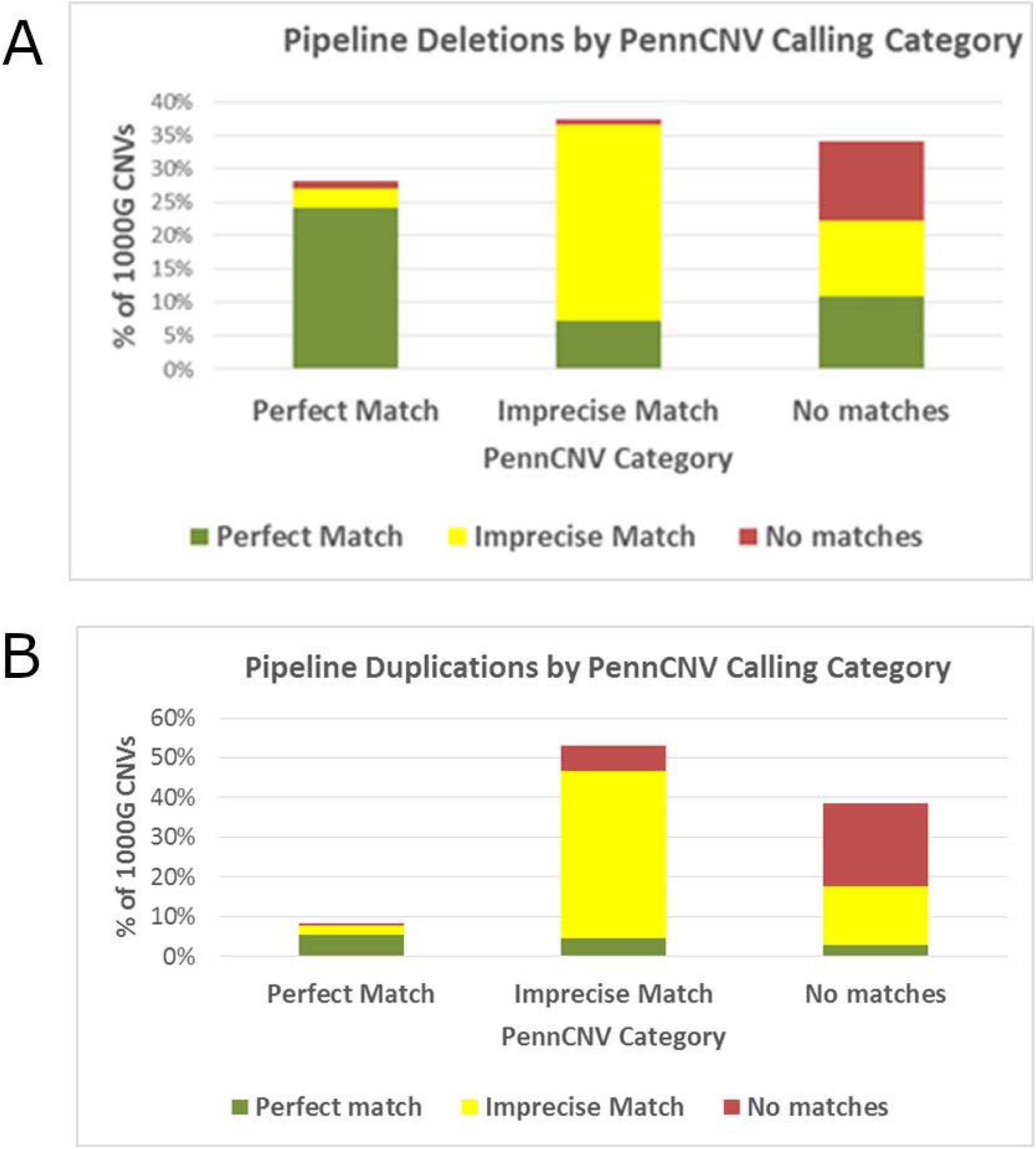
Comparison of sensitivity of CamCNV and PennCNV calling of 1000 Genomes deletions (A) and duplications (B)

**Figure 6.**
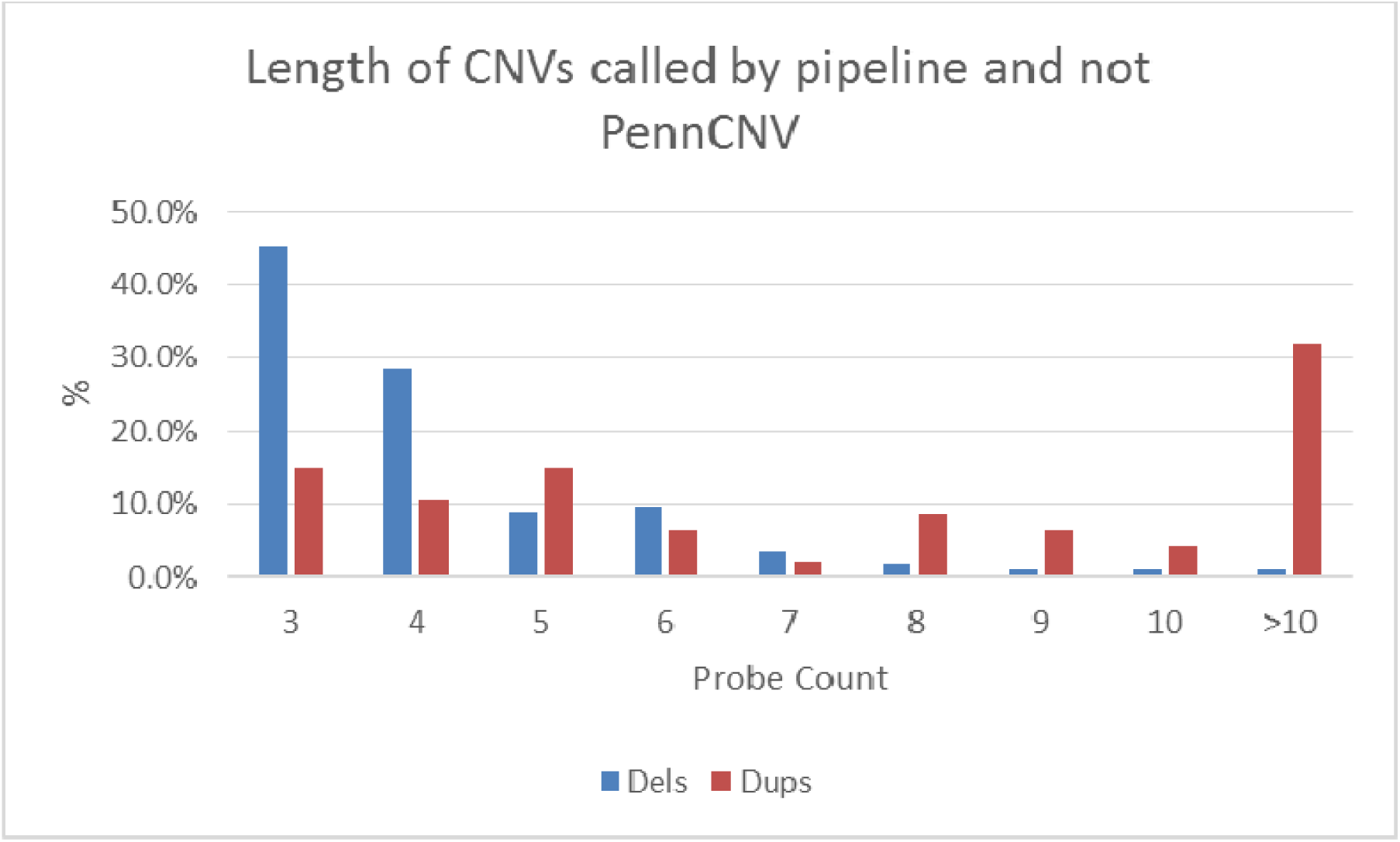
Length of CNVs detected by CamCNV and not by PennCNV

We also estimated the FDR for PennCNV using the BCAC replicate samples. The CamCNV FDR was lower than PennCNV for deletions (Table 8) but higher for short duplications. In the replicate samples CamCNV called around 1.5 times more deletions and ∼1.4 times more duplications than PennCNV, with nearly all the additional deletions having fewer than six probes.

**Table 8.** FDR of CamCNV and PennCNV from replicate samples comparison

| Probes | CamCNV |  | PennCNV |  | CamCNV |  | PennCNV |  |
| --- | --- | --- | --- | --- | --- | --- | --- | --- |
|  | Deletion Count | FDR | Deletion Count | FDR | Duplication Count | FDR | Duplication Count | FDR |
| 3 to 5 | 2872 | 16.2% | 1286 | 26.8% | 1391 | 50.0% | 1097 | 42.6% |
| 6 to 9 | 861 | 1.9% | 825 | 8.2% | 957 | 30.9% | 720 | 18.4% |
| 10+ | 1133 | 1.2% | 1141 | 3.2% | 1679 | 8.5% | 1142 | 11.3% |

## 5 Discussion

CNV calling algorithms often attempt to call rare CNVs together with common CNVs where four or more copy number states will be reflected in multiple overlapping intensity distributions at each probe. The calling problem is simplified when calling rare CNVs as we can assume that nearly all samples will form part of the normal two-copy distribution with a small number of outliers that are potentially one-copy deletions or three-copy duplications. The two-copy intensity distributions vary between probes but the principal component adjustment and conversion of raw intensities to z-scores adjusts for this variance. Thus our method incorporates information from all the samples at each probe; this may explain the improvement in sensitivity compared to the PennCNV method, which looks at each sample individually. From known CNVs we can derive the distribution of z-scores for one-copy deletions and three-copy duplications and calibrate the mean z-score cut-offs for segments to provide an acceptable FDR. From the replicate comparison we observe that calls with more extreme mean z-scores are unlikely to be false positives, so the mean z-score provides an additional metric for the confidence of the calling alongside the number of probes.

A limitation of our method (but also other methods) is that the calling for duplications is less reliable than for deletions, as the duplication intensity distribution has a greater degree of overlap with normal copy samples. Also, there remains the possibility that technical artefacts will cause false calls to cluster at particular probes. For rare variant association tests a proportion of false CNV calls scattered across the chip is not as detrimental as a cluster of false calls at particular probes, so chip-specific probe quality control is an important element of this pipeline. We have tested the method on another Illumina array (iCOGs) and the mean z-score cut-offs used for OncoArray appeared appropriate for that data as well. The cut-offs are likely to be applicable to other Illumina arrays as they are based on the normal distributions of normalised intensity data, but there may be technical reasons for other arrays to have different degrees of overlap between the two-copy and one or three-copy distributions.

## 6 Conclusion

We present a CNV calling pipeline tailored to detect rare CNVs from Illumina genotyping data. For deletions it demonstrates a high degree of sensitivity while controlling the false call rate and is an improvement on one of the most commonly used methods (PennCNV) particularly in detecting deletions covering only a few probes. We believe that this method will be valuable in testing for associations with rare CNVs in large genotyping datasets.

## 7 Declarations

### 7.1 Ethics approval and consent to participate

All participating BCAC studies were approved by their appropriate ethics review board and all subjects provided informed consent.

### 7.2 Consent for publication

Not applicable.

### 7.3 Availability of data and materials

Project name: CamCNV

Project home page: https://github.com/jgd29/CamCNV

Programming language: R

Operating system: Platform independent

Example code and a test dataset with 1000 Genomes OncoArray genotyping is available at the project home page.

The BCAC OncoArray dataset supporting the conclusions of this article is available in the dbGap repository, Study ID: phs001265.v1.p1 https://www.ncbi.nlm.nih.gov/projects/gap/cgi-bin/study.cgi?study_id=phs001265.v1.p1

The 1000 Genomes Phase 3 structural calls are available from ftp://ftp-trace.ncbi.nih.gov/1000genomes/ftp/phase3/integrated_sv_map/ALL.wgs.integrated_sv_map_v2.20130502.svs.genotypes.vcf.gz

### 7.6 Authors’ contributions

JD and DFE designed the project. JD, DFE, LW and JT developed the pipeline. KM performed the imputation. All authors read and approved the final manuscript.

## 7.7 Acknowledgements

The authors thank all BCAC investigators for providing access to the BCAC dataset. The BCAC studies are listed in supplementary table 2. We thank all individuals who took part in these studies and all the researchers, clinicians, technicians and administrative staff who have enabled this work to be carried out.

---

**Supplementary Table 1.** Overlap of called CNVs with known CNV regions (Map from Zarrei et al.)

| Category | Probes | Zarrei Stringent Map | % covered | Zarrei Less Stringent Map | % covered |
| --- | --- | --- | --- | --- | --- |
| All probes on chip | 533,631 | 20970 | 3.9% | 43233 | 8.1% |
| Probes in CNV calling | 480,982 | 13618 | 2.8% | 33237 | 6.9% |
| Deletions <0.5% | 306,947 | 8159 | 2.7% | 21540 | 7.0% |
| Deletions 0.5% to 1% | 542 | 163 | 30.1% | 296 | 54.6% |
| Deletions >1% | 186 | 94 | 50.5% | 125 | 67.2% |
| Duplications <0.5% | 382,723 | 8660 | 2.3% | 24881 | 6.5% |
| Duplications 0.5% to 1% | 997 | 333 | 33.4% | 458 | 45.9% |
| Duplications >1% | 768 | 187 | 24.3% | 231 | 30.1% |

**Supplementary Table 2.** BCAC Studies

| Acronym | Study | Institution | Country |
| --- | --- | --- | --- |
| ABCFS | Australian Breast Cancer Family Study | University of Melbourne | Australia |
| ABCS | Amsterdam Breast Cancer Study | Netherlands Cancer Institute (NKI) | Netherlands |
| ABCTB | Australian Breast Cancer Tissue Bank | University of Sydney | Australia |
| AHS | Agricultural Health Study | National Cancer Institute (NCI) | USA |
| BBCC | Bavarian Breast Cancer Cases and Controls | University Hospital Erlangen | Germany |
| BBCS | British Breast Cancer Study | London School of Hygiene and Tropical Medicine | UK |
| BCEES | Breast Cancer Employment and Environment Study | University of Western Australia | Australia |
| BCINIS | Breast Cancer In Northern Israel Study | National Israeli Cancer Control Center (NICCC) | Israel |
| BREOGA N | Breast Oncology Galicia Network | Galician Foundation of Genomic Medicine/SERGAS | Spain |
| BSUCH | Breast Cancer Study of the University of Heidelberg | University of Heidelberg | Germany |
| CBCS | Canadian Breast Cancer Study | British Columbia Cancer Agency | Canada |
| CCGP | Crete Cancer Genetics Program | University Hospital of Heraklion | Greece |
| CECILE | CECILE Breast Cancer Study | Inserm (National Institute of Health and Medical Research) | France |
| CGPS | Copenhagen General Population Study | Copenhagen University Hospital | Denmark |
| CPSII | Cancer Prevention Study-II Nutrition Cohort | American Cancer Society (ACS) | USA |
| CTS | California Teachers Study | Beckman Research Institute of City of Hope | USA |
| EPIC | European Prospective Investigation Into Cancer and Nutrition (BPC3) | The International Agency for Research on Cancer (IARC) | Various |
| ESTHER | ESTHER Breast Cancer Study | German Cancer Research Center (DKFZ) | Germany |
| FHRISK | Family History Risk Study | University of Manchester | UK |
| GC-HBOC | German Consortium for Hereditary Breast & Ovarian Cancer | University Hospital of Cologne | Germany |
| GENICA | Gene Environment Interaction and Breast Cancer in Germany | Dr. Margarete Fischer-Bosch-Institute of Clinical Pharmacology Stuttgart | Germany |
| GESBC | Genetic Epidemiology Study of Breast Cancer by Age 50 | German Cancer Research Center (DKFZ) | Germany |
| HABCS | Hannover Breast Cancer Study | Hannover Medical School | Germany |
| HEBCS | Helsinki Breast Cancer Study | Helsinki University Central Hospital | Finland |
| HMBCS | Hannover-Minsk Breast Cancer Study | Hannover Medical School | Belarus |
| HUBCS | Hannover-Ufa Breast Cancer Study | Institute of Biochemistry and Ufa Scientific Center of Russian Academy of Sciences (UCRAS) | Russia |
| KARMA | Karolinska Mammography Project for Risk Prediction of Breast Cancer – Cohort Study | Karolinska Institutet | Sweden |
| KBCP | Kuopio Breast Cancer Project | University of Eastern Finland | Finland |
| LMBC | Leuven Multidisciplinary Breast Centre | Flanders Institute for Biotechnology (VIB) | Belgium |
| MaBCS | Macedonian Breast Cancer Study | Macedonian Academy of Sciences and Arts (MASA) | Republic of North Macedonia |
| MARIE | Mammary Carcinoma Risk Factor Investigation | German Cancer Research Center (DKFZ) | Germany |
| MBCSG | Milan Breast Cancer Study Group | Fondazione IRCCS Istituto Nazionale dei Tumori (INT) and the Istituto Europeo di Oncologia (IEO) | Italy |
| MCBCS | Mayo Clinic Breast Cancer Study | Mayo Clinic | USA |
| MCCS | Melbourne Collaborative Cohort Study | Cancer Council Victoria | Australia |
| MEC | Multiethnic Cohort | University of Southern California | USA |
| MISS | Melanoma Inquiry of Southern Sweden | Lund University | Sweden |
| MMHS | Mayo Mammography Health Study | Mayo Clinic | USA |
| MTLGEB<br>CS | Montreal Gene-Environment<br>Breast Cancer Study | McGill University | Canada |
| NC-BCFR | Northern California Breast<br>Cancer Family Registry | Cancer Prevention Institute of<br>California | USA |
| NCBCS | North Carolina Breast Cancer<br>study | University of North Carolina at<br>Chapel Hill | USA |
| NHS | Nurses' Health Study | Harvard School of Public Health<br>(HSPH) | USA |
| NHS2 | Nurses' Health Study 2 | Harvard School of Public Health<br>(HSPH) | USA |
| OFBCR | Ontario Familial Breast Cancer<br>Registry | Mount Sinai Hospital | Canada |
| ORIGO | Leiden University Medical<br>Centre Breast Cancer Study | Leiden University Medical Centre | Netherlands |
| PBCS | NCI Polish Breast Cancer Study | National Cancer Institute (NCI) | Poland |
| PLCO | The Prostate, Lung, Colorectal<br>and Ovarian (PLCO) Cancer<br>Screening Trial | National Cancer Institute (NCI) | USA |
| RBCS | Rotterdam Breast Cancer Study | Erasmus University Medical Center<br>Rotterdam | Netherlands |
| SEARCH | Study of Epidemiology and Risk<br>factors in Cancer Heredity | University of Cambridge | UK |
| SISTER | The Sister Study | National Institute of Environmental<br>Health Sciences (NIEHS) | USA |
| SMC | Swedish Mammography Cohort | Karolinska Institutet | Sweden |
| SZBCS | IHCC-Szczecin Breast Cancer<br>Study | Pomeranian Medical University | Poland |
| UCIBCS | UCI Breast Cancer Study | University of California Irvine | USA |
| UKBGS | UK Breakthrough Generations<br>Study | Institute of Cancer Research (ICR) | UK |
| USRT | US Radiologic Technologists<br>Study | National Cancer Institute (NCI) | USA |

**Supplementary Figure 1.**
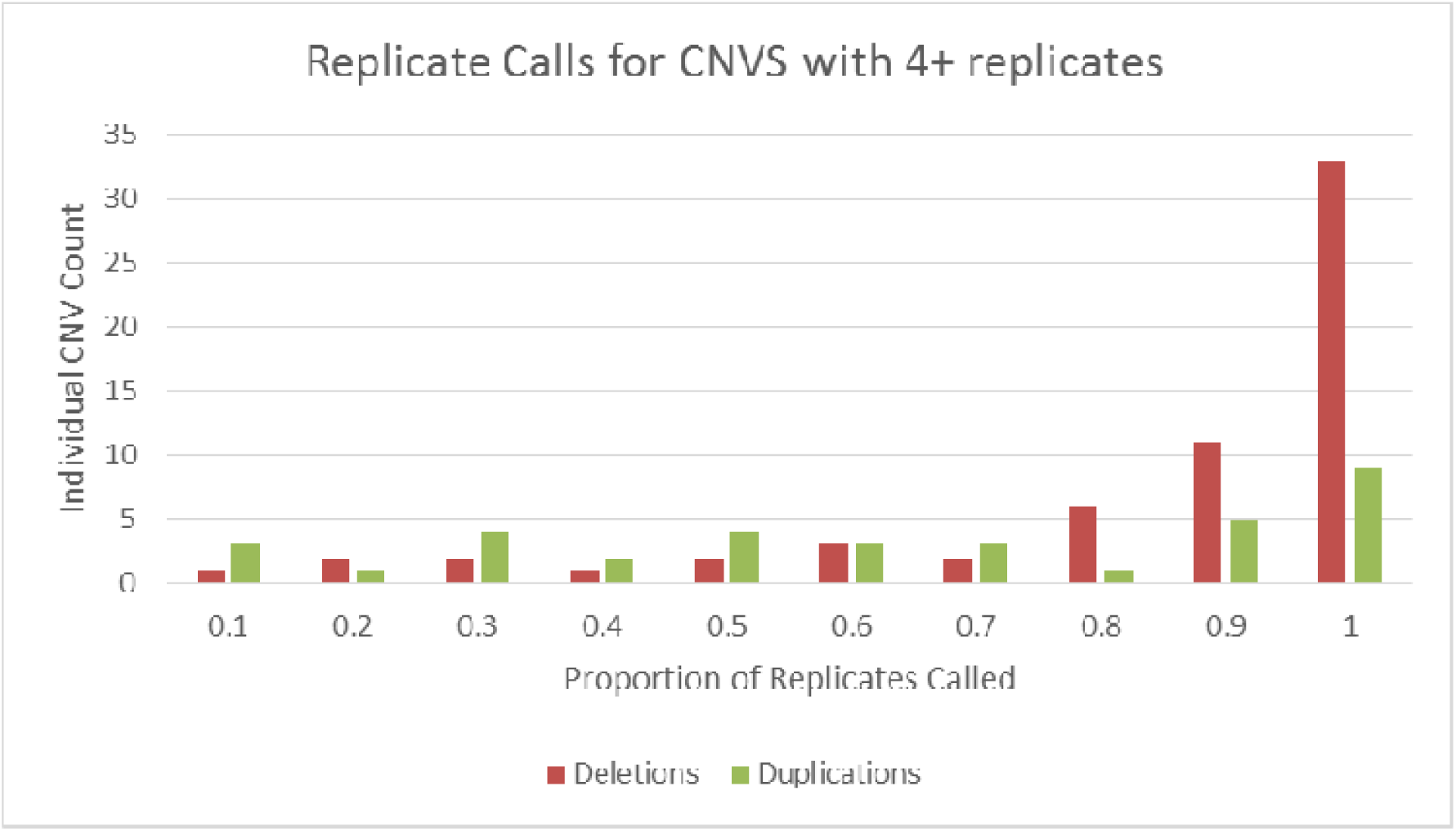
Replication rate for 1000 Genomes called CNVs with more than three replicates

**Supplementary Figure 2.**
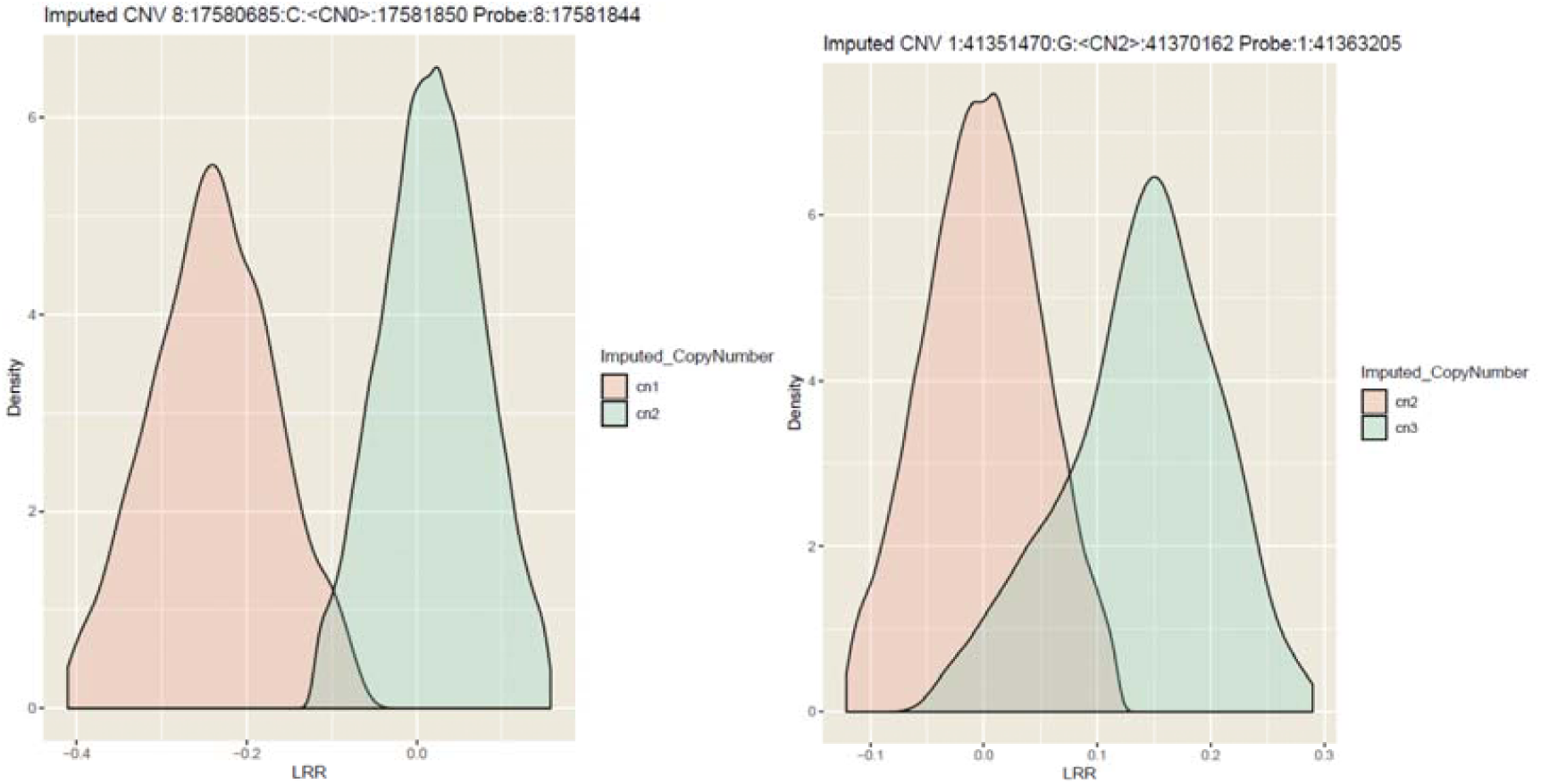
Examples of distributions of intensities by imputed copy number state for a single probe in a common deletion and a probe in a duplication.

